# Contribution of livestock marketing chains and role played by stakeholders’ knowledge, attitude and practice in spreading cystic hydatidosis to Busia Town, Kenya, 2018

**DOI:** 10.1101/638502

**Authors:** Henry Joash Ogutu, Maurice Owiny, Bernard Bett, Christina Otieno

**Affiliations:** Kenya Field Epidemiology and Laboratory Training Program, Nairobi, Kenya; School of Public Health, Moi University, Eldoret, Kenya; International Livestock Research Institute, Nairobi, Kenya

**Author notes:** **Corresponding author:** (HJO).

**Keywords:** Cystic hydatidosis, cattle marketing chains, KAP, Busia-Kenya

## Abstract

**Background:** Cystic hydatidosis (CH), a neglected parasitic zoonosis, is endemic in many parts of Kenya and could be spread along livestock marketing chains. Poor knowledge, attitude and practices (KAP) enables this spread in remote areas with inadequate public health services. We estimated prevalence, identified possible origin of CH to Busia, Kenya and assessed KAP among cattle owners and abattoir workers.

**Methods and Principal Findings:** We conducted a cross-sectional study on slaughtered livestock and interviewed their owners and abattoir workers in Busia in May–June 2018. We used visual observation, palpation and incision to identify cysts. Polymerase chain reaction (PCR) was used for confirmatory diagnosis. Epi Info 7 was used to calculate descriptive and associative statistics. Of 302 carcasses inspected, cysts were visualized in nine (2.98%, 95% Confidence Interval (CI): 1.46–5.78). Fourteen samples were collected and 13 (92.86%) were positive on PCR (sensitivity=92%, specificity=95%). All carcasses with cysts were from West Pokot County, which borders Busia to the north. We interviewed 310 participants: 260 were males (83.87%, 95% CI: 79.19 – 87.69); median age was 41 years (range=21-69). Dogs were kept by 221 (71.99%, 95% CI: 66.55 – 76.87), of which 83 (37.56%, 95% CI: 28.33 – 48.52) improperly disposed of dog faeces. Home slaughtering was practiced by 196 (63.23%, 95% CI: 58.78-69.80), of which 115 (58.67%, 95% CI: 51.44-65.64) were not inspected and 85 (43.37%, 95% CI: 36.32-50.62) fed raw organs to dogs. Adequate knowledge was associated with butcher ownership (P-value = 0.002), age ≥35 years (P-value = 0.002) and higher literacy level (P-value <0.001).

**Conclusions and Significance:** There is non-negligible risk of CH in Busia communities which might worsen with time given that the county is connected to endemic areas through livestock trade. Poor KAP by the people on the disease calls for need to implement information, education and communication campaigns to improve KAP on CH in the area.

**Author summary:** Cystic hydatidosis is a globally neglected parasitic zoonosis which is endemic in many parts of the world including Kenya. It is majorly a problem among pastoral communities where there is close contact between human, livestock and dogs. Busia County, in Western Kenya is part of a livestock marketing chain between Kenya and Uganda. Animals from high endemic regions in Uganda and Kenya can easily spread the parasite to Busia through improper disposal of their infested organs. Non-pastoral communities like Busia may not have much cumulative experience about the disease though their practices may contribute to the perpetuation of the parasite in their environment. The parasite is gradually spreading to new areas and it is very important to the public health players in Kenya to take action so as to prevent further spread of this disease. Findings from this study show that the disease is no longer limited to pastoral communities only. There is need for the implementation of information, education and communication campaigns to improve the knowledge, attitude and practices of Busia community and other non-endemic regions on the disease.

## Introduction

Hydatidosis is a neglected parasitic zoonotic disease caused by larval stage of *Echinococcus* and affects mostly dogs, livestock and humans [1,2]. The parasite has four species, but only two are of public health importance; *E. granulosus* which causes cystic hydatidosis (CH), commonly occurs in tropical regions and infests ungulates as its prime intermediate host hence its significance in livestock and *E. multilocularis*, which causes alveolar hydatidosis (AH) and occurs in the temperate regions [3]. Others are *E. vogeli*, and *E. oligarthrus* [3,4]. Hydatidosis has been reported in Europe and south eastern Australia. It is endemic in China, Indian Subcontinent and Middle East and re-emerging in the former Soviet Republics. In Africa *E. granulosus* is a particular problem in Northern and Eastern Africa countries including Kenya [2].

Hydatidosis affects 2–3 million people worldwide in extensive livestock farming areas. In 2014 the WHO estimated that it caused more than 3,000 human and animal deaths. These deaths contributed to economic losses estimated at three billion United States Dollars (USD) covering costs on interventions, livestock organ condemnation and reduced livestock productivity. Livestock related losses in Kenya are estimated at more than 240,000 USD annually [5].

Busia, a border town in Kenya, is a major livestock market for traders in Kenya and Uganda. There are fears that the parasite is being introduced into the county via livestock marketing chains by improper disposal of infested organs [5]. Previous studies on the disease have put less focus on the risks of spread via livestock trade that connect high and low endemic areas. Findings from such studies could aid public health actors in formulating evidence-based prevention and control policies for this disease. This study estimated prevalence of CH in cattle slaughtered at Busia abattoirs, identified possible origin of CH to Busia and assessed knowledge, attitude and practices (KAP) on hydatidosis among livestock owners and people working in slaughter houses in Busia, Kenya.

## Methods

### Study design

We conducted a cross-sectional study on cattle carcasses and interviewed abattoir workers as well as people who presented their cattle for slaughter on their knowledge attitude and practices on hydatidosis. The study was conducted in two busy abattoirs in Busia Town; Busia Municipal and Amerikwai abattoirs form May to June 2018. Busia Town is at the border of Kenya and Uganda in Busia County (Figure 1). It has a total human population of 111,345 and livestock population of 215,871 out of which cattle are 132,804 [6]. The main economic activity in Busia town is trade with neighboring Uganda, but crop and livestock farming is also done in small scale in the outskirts of the town. We included male and female cattle of all age categories destined for slaughter at the two abattoirs. All carcasses, whose owners consented, were eligible for inspection. Consenting adult cattle owners and abattoir workers were assessed on their KAP on hydatidosis. Carcasses condemned for other infections like emphysema, liver flukes and tuberculosis were not eligible for consideration into the study. Cattle whose owners could not be reached or traced for interviewing were not eligible.

**Fig 1:**
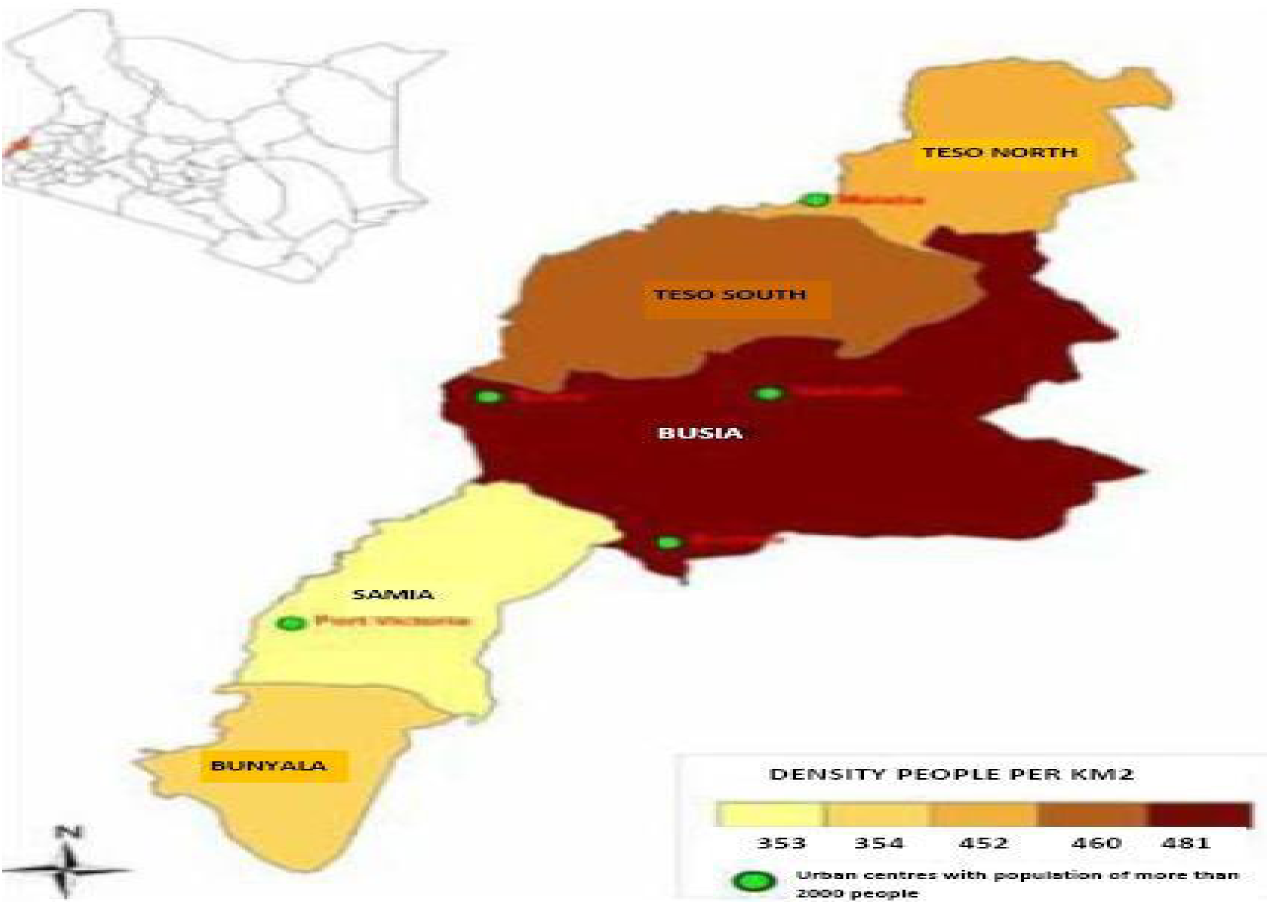
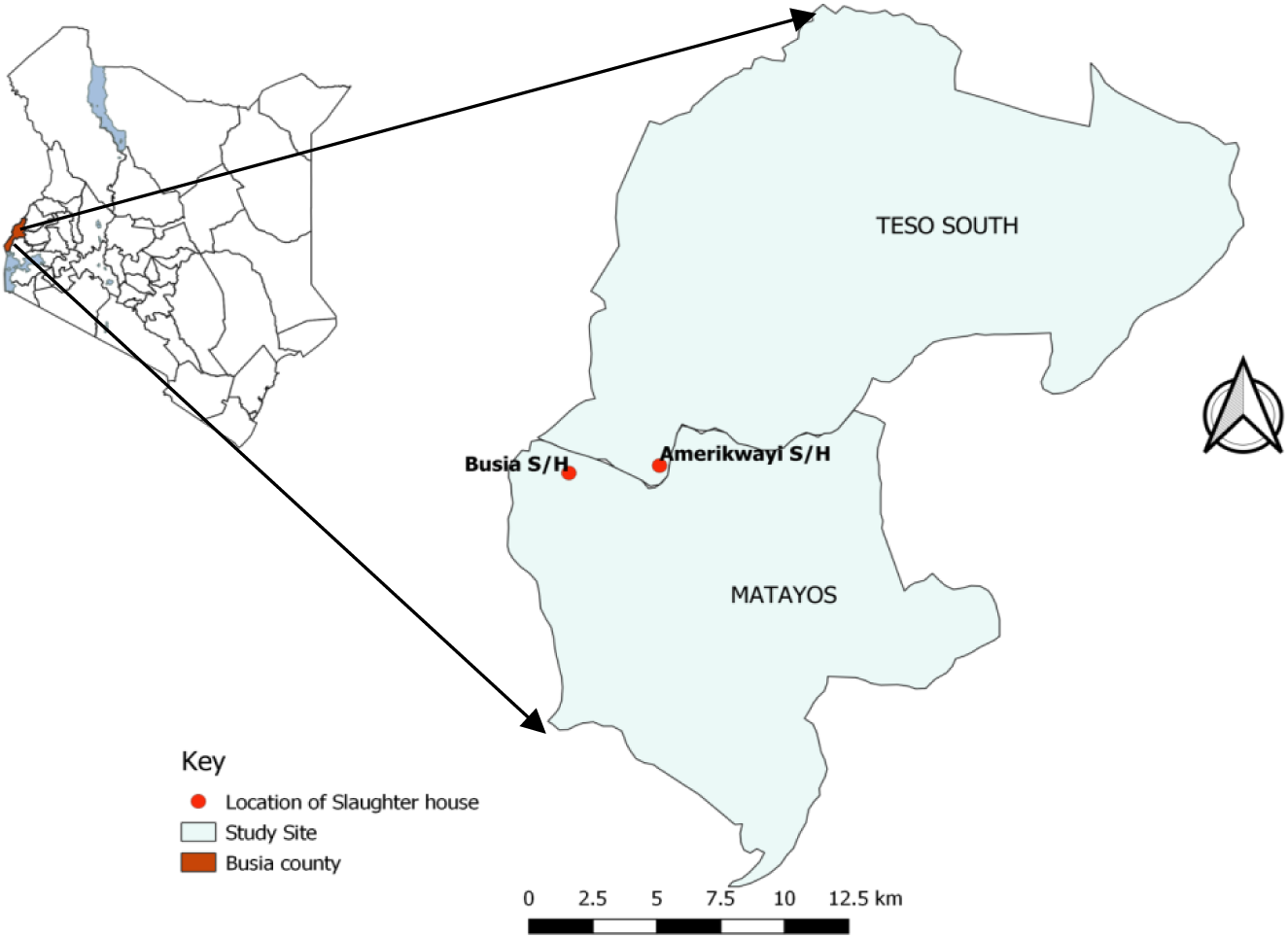
Map of study area. Brown rectangle in Kenyan map represent the location of Busia in relation to neighboring Uganda, Red circle represent location of Busia Municipal and Amerikwai abattoirs. *This figure was created for this manuscript in QGIS using open source data from ESRI and GPS points collected during data collection in the field.*

### Sample size estimation and assumptions

A minimum sample size of 294 was calculated using the Cochrane formula with the following assumptions; *a priori* prevalence of 25.8 % (1), 95 % as the level of confidence, 5% desired absolute precision and 10% anticipated non-response rate. The sample size was proportionately divided between the two abattoirs based on average monthly slaughter figures as per the monthly meat hygiene reports over a period of three years.

### Variables and measurements

Among the variables collected at animal level included sex, estimated age, breed and origin of each animal. We used questionnaires to collect data on whether or not the respondent had gone through training on CH, knowledge on the mode of transmission and control measures for the disease, importance of deworming dogs and livestock to control hydatidosis, slaughtering animals at home, keeping dogs at home, feeding dogs on raw meat, disposal of infected organs and dog feces at home.

### Data collection

We conducted ante-mortem examinations of all study cattle to estimate age by dentition [7,8], identify breed, determine sex and record the origin of each animal. The origin of each animal was confirmed from the animal movement permits and ‘no objection’ forms filed in the Busia veterinary offices and also from interviews held with livestock traders and farmers who brought their animals for slaughter. Post-mortem inspections were conducted immediately after slaughtering, skinning and evisceration. Standard morphological meat inspection procedures including visual observation, palpation and systematic incision by making deep longitudinal cuts in organs and muscles. according to Kenya Meat Control Act, CAP 356, were used to determine infestation status of each carcass. Number of cysts per organ were counted and recorded and cysts were removed whole and put in zipped polythene bags in cool boxes with icepacks for conventional polymerase chain reaction (PCR) tests at the International Livestock Research Institute (ILRI) field laboratory in Busia. Each sample was appropriately labelled for ease of traceability in the laboratory.

### Laboratory methods

The samples were gradually frozen to −20°C, kept for one month then processed and stored in 70% ethanol. For DNA extraction, up to 20 mg of tissue samples was excised and placed in a nuclease-free microfuge tube. We added 300 microliters (μL) of digestion buffer A to the tissue and 12 μL of proteinase K and left to incubate at 55°C for 1.5 hours. We then added 300 μL of buffer SK to the lysate and mixed by vortexing and then added 300 μL of 100% ethanol. A micro spin column with a provided collection tube was assembled and up to 600 μL of the mixture was applied to the spin column assembly. The unit was capped and centrifuged for three minutes at 8,000 rotations per minute (RPM). After centrifugation, we discarded the flow-through and reassembled the spin column with its collection tube. This was repeated until all the lysate had passed through the column. To wash the bound DNA, we applied 500 μL of wash solution A to the column and centrifuged the unit for one minute at 14000 RPM. After centrifugation, we discarded the flow-through and reassembled the spin column with its collection tube. We applied 500 μL of wash solution A to the column and centrifuged the unit for two minutes at 14000RPM. The spin column was detached from the collection tube and discarded the collection tube and flow-through. We assembled the spin column with DNA bound to the resin with a provided 1.7 mL elution tube. Two hundred microliters of Elution Buffer B was added to the center of the resin bed then allowed to stand for 10 minutes. It was then centrifuged for one minute at 6000RPM. A portion of Elution Buffer B passed through the column which allowed for the hydration of the DNA to occur. We again centrifuged at 14000RPM for an additional two minutes to collect the total elution volume. The purified genomic DNA was stored at −20°C for one day to await PCR process.

#### Identification of *nad5* gene

Conventional PCR was carried out on all the 14 hydatid cysts DNA isolated. The PCR primers Mit-F/Mit-R were used to amplify a 562 bp fragment of the mitochondrion the NADH-Ubiquinone oxidoreductase (complex I), chain 5 N-terminus (*nad5* gene) of *E. granulosus* (Gen Bank accession No. ARO49807). The PCR amplification reactions containing 3 μL mtDNA, 0.5 μL each of the forward and reverse primers (this study), and 12.5 μL of *Taq* PCR Master Mix (Qiagen) in a final reaction volume of 28 μL. After denaturation at 95°C for 10 min, amplification cycles were performed for four-stage, 25 cycles of 95°C for 30 seconds (s), 58°C for 30 s, and 72°C for 30 s for seven cycles in stage one, 95°C for 30 s, 56°C for 30 s, and 72°C for 30 s for seven cycles in stage two, 95°C for 30 s, 55°C for 30 s, and 72°C for 30 s for seven cycles in stage three, 95°C for 30 s, 54°C for 30 s, and 72°C for 30 s for four cycles in stage four, followed by 72°C for 10 min and cooling to 10°C. PCR products were loaded on 1.2% (w/v) Hi-Standard Agarose gel (AGTC Bio-products Limited, Hessle, UK) in 1X Tris-Boric-EDTA and stained with 0.5 μg/ml Safe White Nucleic Acid Stain (NBS Biologicals, Cambridge-shire, UK). Electrophoresis was carried out for 40 min at 190 V. The bands were visualized in UV trans-illuminator and digitally photographed.

### Knowledge, attitude and practices

We used a pre-tested questionnaire with open ended and closed-questions for interviews. The interviews were conducted in a separate room within the abattoir compounds to maintain confidentiality.

### Statistical methods

The collected data were entered, cleaned and analyzed using Epi Info™ 7.1.4.0 (CDC, Atlanta, GA, USA). Measures of central tendency and dispersion for continuous variables and frequencies, proportions and 95% CI for categorical variables were calculated. We assigned scores to knowledge questions in the questionnaire. A correct response earned a score of one (1) while an incorrect or *“I don’t know”* response scored a zero (0). Adequate knowledge was considered as a total score above or equal to half (≥5) of the overall score (10). We used bivariate and logistic regression to examine the factors associated with adequate knowledge among the study participants. From the bivariate analysis, variables that had P-values of ≤ 0.1 were entered into a multivariate regression model. The final model was arrived at using backward stepwise elimination method where variables with P-values ≤ 0.05 were considered to have statistical associations with adequate knowledge, as dependent coefficient. The participants who considered keeping dogs and livestock in same homestead to increase the risk of hydatidosis were considered to have a good attitude. Regular deworming of livestock and dogs, good meat hygiene and proper disposal of infected organs and dog faeces were considered as good practices.

### Ethical issues

We obtained written permission from Busia County Director of Veterinary Services (CDVS) to access the abattoirs and sought verbal consent from abattoir managers to access abattoir compounds. Participation by cattle owners was sought through signing informed consent. Ethical approval for this study was obtained from institutional research and ethics committee (IREC) of Moi University (IREC No. 0001850).

## Results

### Descriptive statistics

In the study, a total of 302 cattle carcasses were inspected. Of these, 188 (62.25%) originated from Busia County while others originated from Uganda and other parts of Kenya (Figure 2). Local breeds of cattle comprised 295 (97.68%, 95% Confidence Interval (CI): 95.08–98.98). The inspected carcasses consisted of 222 (73.51%, 95% CI: 68.09 – 78.32) male cattle.

**Fig 2:**
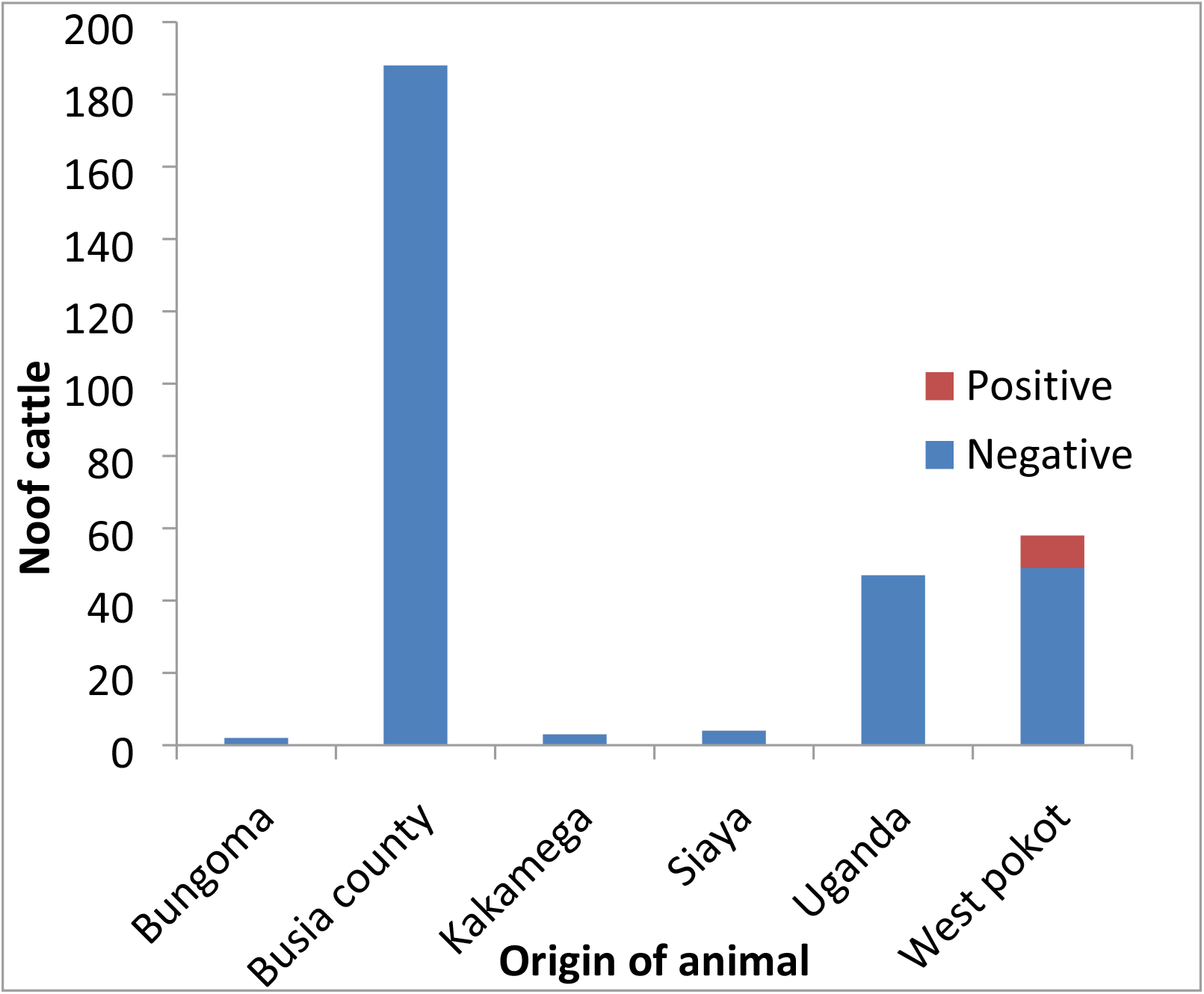
Distribution of source of cattle by county and their infection status, Busia town abattoirs 2018 (n=302).

Majority of the carcasses inspected, 144 (47.68%) were aged between 4–6 years and 18 (5.96%, 95% CI: 3.67–9.42) were aged above nine years. Hydatid cysts were visualized in nine (2.98%) of the inspected carcasses. Among these, eight were female (Table 1).

Out of the nine carcasses with cysts, five (55.56%) had multiple organ infestations. The main infested organs were liver (n=7) and lung (n=4). The total number of samples collected for PCR confirmation was 14 out of which 13 (92.86%) samples turned positive on PCR test (sensitivity = 92% and specificity = 95%). From the laboratory results, the samples whose bands were visualized in ultra violet (UV) trans-illuminator were interpreted as positive on the PCR test. One sample calcified during preservation and therefore the DNA could not be extracted from it for PCR testing. All the positive carcasses were animals from West Pokot County.

**Table 1:** Proportions of cattle by age, gender and infection status at Busia town abattoirs 2018 (n=302)

| Age grp | Male |  | Total male | Female |  | Total female | Total carcasses |
| --- | --- | --- | --- | --- | --- | --- | --- |
|  | Positive | Negative |  | Positive | Negative |  |  |
| 1-3 | 0 | 13 | 13 | 0 | 5 | 5 | **18** |
| 4-6 | 0 | 101 | 101 | 2 | 41 | 43 | **144** |
| 7-9 | 0 | 93 | 93 | 6 | 23 | 29 | **122** |
| >9 | 1 | 14 | 15 | 0 | 3 | 3 | **18** |

### Knowledge, attitude and practices

We interviewed 310 cattle owners, of whom 260 (83.87%, 95% CI: 79.19 – 87.69) were male. The overall median age was 41 years (range 21-69 years). Livestock farmers comprised 158 (50.97%, 95% CI: 45.27 – 56.65) of the study participants (Table 2). When asked about their level of education, 116 (37.42%, 95% CI: 32.06 – 43.06) said that they had completed primary education, 92 (29.67%, 95% CI: 24.71 – 35.15) had completed secondary education and 58 (18.71%, 95% CI: 14.62 – 23.60) did not have any formal education (Table 2).

**Table 2:** Literacy level of participants and their role in the value chain, Busia town abattoirs, 2018.

| Factor | Frequency | Percentage |
| --- | --- | --- |
| **Level of education (n=310)** |  |  |
| None | 58 | 18.70 |
| Primary completed | 116 | 37.42 |
| Secondary completed | 92 | 29.68 |
| Tertiary completed | 44 | 14.20 |
| **Role in value chain (n=310)** |  |  |
| Livestock farmer | 158 | 50.97 |
| Butchery owner | 65 | 20.97 |
| Livestock trader | 39 | 12.58 |
| Butchery attendant | 21 | 6.77 |
| Butcher man/flayer | 17 | 5.48 |
| Meat inspector | 5 | 1.61 |
| Intern/student | 3 | 0.97 |
| Abattoir cleaner | 2 | 0.65 |

On knowledge of cystic hydatidosis, 197 (63.55%, 95% CI: 57.89 – 68.86) participants said that they had heard about the disease. The participants who knew the role of dogs in transmission of the disease were 53 (17.10%, 95% CI: 13.37 – 23.84). The effects of hydatidosis on livestock were known to 175 (56.45%) of the respondents, out of which 161 (92.00%) were butchery owners. On average, the participants who scored ≥5 were 40 (12.90%).

When we assessed the attitude of the participants on the disease, 162 (54.00%) disagreed that there is a risk of hydatid disease transmission to livestock or humans by having a dog on the same compound with livestock; 91 (30.33%) agreed, 32 (10.67%) strongly agreed and 15 (5%) strongly disagreed to the question. Among the participants who answered the question regarding the importance of deworming dogs to control the disease, 130 (43.33%) disagreed, 119 (39.67%) agreed, 43 (14.33%) strongly agreed and eight (2.67%) strongly disagreed. The participants who disagreed that disposing or condemning infected organs was a waste of food were 159 (51.96%), while 95 (31.05%) strongly disagreed, 42 (13.73%) agreed, and 10 (3.27%) strongly agreed. Those who agreed that keeping their livestock dewormed and clean was a reflection of their social status were 177 (57.65%), while 88 (28.66%) strongly agreed, 42 (13.68%) disagreed, and none strongly disagreed. A good attitude towards the disease was held by 123 (39.68%).

Assessment of study participants on their practice on the disease revealed that 256 (85.62%, 95% CI: 81.12 – 89.39) dewormed their livestock and 124 (48.44%, 95% CI: 42.17 – 54.74) of them dewormed their livestock after every 3 months. Dogs were kept at home by 221 (71.29%, 95% CI: 66.55 – 76.87) participants. Among the dog keepers, 93 (42.08%, 95% CI: 35.49 – 48.89) dewormed their dogs and 37 (39.78%, 95% CI: 29.78 – 50.46) dewormed at an interval of three months. Methods of disposing dog faeces included burying, 95 (42.99%, 95% CI: 36.37 – 49.80), doing nothing and open disposal, 83 (37.56%, 95% CI: 28.33 – 48.52). Feeding their dogs on raw meat was admitted by 120 (54.30%, 95% CI: 47.48 – 61.00) dog keepers, while 196 (63.23%, 95% CI: 58.78 – 69.80) cattle owners admitted that they sometimes slaughtered animals at home. However, 115 (58.67%, 95% CI: 51.44 – 65.64) of the meat slaughtered at home was not inspected by qualified meat inspectors and 85 (43.37%, 95% CI: 36.32 – 50.62) of raw organs of animals slaughtered at home were fed to dogs. At bivariate analysis, religion (*P*<0.0447), gender (*P*<0.0354), age (*P*<0.0046), education (0.0012) and occupation (0.0085) had a statistical association with adequate knowledge (Table 3).

**Table 3:** Bivariate analysis with knowledge as a coefficient of other variables.

| Variable | OR (95% CI) | P value |
| --- | --- | --- |
| Age (> 35 years) | 88.44 (51.79 – 945.74) | 0.0046 |
| Education (primary or less) | 56.13 (40.33 – 194.67) | 0.0012 |
| Gender | 83.87 (0.87 – 112.44) | 0.0354 |
| Marital status (single) | 8.71 (2.45 – 44.12) | 6.5437 |
| Occupation (butchery owner) | 51.94 (33.56 – 98.89) | 0.0085 |
| Religion (none) | 0.97 (0.56 – 2.44) | 0.0447 |

**OR: Odds Ratio, CI: Confidence Interval**

On multivariate analysis, occupation (being a butchery owner) (*P*<0.002), age above 35 years (*P*<0.002) and literacy level (*P*<0.001) were independently associated with adequate knowledge on CH (Table 4).

**Table 4:** Multivariate logistic regression with knowledge as random effect variable to age, level of education and occupation.

| Coefficient | AOR (95% CI) | Standard error | P value |
| --- | --- | --- | --- |
| Age (> 35 years) | 73.43 (21.71 – 1342.72) | 1.0216 | 0.002 |
| Education (primary or less) | 56.13 (40.33 – 194.67) | 0.27990 | 0.001 |
| Gender | 83.87 (0.87 – 112.44) | 1.0293 | 1.581 |
| Marital status (single) | 8.71 (2.45 – 44.12) | 1.0319 | 4.056 |
| Occupation(butchery owner) | 0.47 (0.12 – 0.86) | 0.4361 | 0.002 |
| Religion (none) | 0.97 (0.56 – 2.44) | 0.4361 | 0.223 |

*AOR: Adjusted Odds Ration, CI: Confidence Interval*

## Discussion

The study found that there is a risk of cystic hydatidosis spreading to Busia town via cattle trade from endemic areas like West Pokot County. Some of these cattle slaughtered in Busia Town originate from Uganda and other parts of Kenya that are infested with the parasite. In Busia, the parasite may spread further through improper disposal of infested organs. The study identified gaps in the participants’ knowledge, attitude and practice regarding awareness on CH, risks of keeping dogs on same compound with livestock, practicing poor meat hygiene, improper disposal of dog faeces and infested organs of animals slaughtered at home in absence of qualified meat inspectors and not deworming of their dogs and livestock regularly. These gaps on KAP can increase the risk of infestation with the parasite.

Most of the cattle slaughtered at Busia town abattoirs during the study period came from within Busia County. Male cattle formed higher percentage of the cattle slaughter figures than females because they grow faster are heavier in weight and attain their mature weight earlier than females. Thus males give higher returns in profit after sale of their carcasses as butchery owners buy cattle for slaughter based on weight and body size. The local breeds of cattle (zebu), being the majority of cattle in Busia County [6], have slow growth rate and get their optimum weight at an average age of four years [9]. This could explains why most of the slaughtered cattle at the two abattoirs were between 4-6 years old.

The reported prevalence of 3% in this study shows the extent to which the infestation can easily spread over time from its known endemic areas. The possibility of spread is very important to the public health stakeholders in the country to take action so as to avoid the spread of this disease to non-endemic regions in Busia [5,10]. The trend of spread over time should be a warning to Kenya’s public health players that the disease is no longer a problem to pastoral communities only. The study findings confirmed that the infestation rates of cattle increased significantly with age [11,12]. This study revealed that liver was the main infested tissue; a similar finding was seen in studies done in Kermanshah province, west of Iran and in slaughter houses in Maasailand and Turkana in Kenya [10,13]. Other studies have found that lungs are the main infested tissues [11,12,14]. Other findings in this study are that female cattle are more likely to be infested with CH than males, similar to a study done in Tabriz area, Northwest of Iran [12], a study done in Central Ethiopia [11] and a study done in Libya [15]. The higher prevalence in female cattle may be correlated to the fact that the females are kept for reproductive purpose hence they live for longer periods while most male cattle are slaughtered at an early age. However, both male and female are at risk of contracting the disease [5].

The findings of the KAP survey showed that the beef value chain in the study area was dominated by men, who have culturally been cattle traders and they tend to dominate livelihood activities that generate financial income as was seen in studies done in Uganda [16] and Pakistan [7]. Most of the participants, especially farmers, did not have much knowledge about hydatidosis. Among those who knew the effects of CH, the majority were butchery owners as they were more familiar with the direct losses due to condemnation of infested organs and carcasses [7]. Results from this study showed that non-pastoral communities like those found in Busia Kenya are unfamiliar with CH. Participants above the age of 35 years were more aware of the disease than younger people. The statistical association between age and knowledge on hydatid disease could be due to the cumulative experience and insights about the disease that accrues with age [17].”

A large number of the participants disagreed that there is a risk of transmission of the disease by keeping a dog on the same compound with the livestock. A majority of them also did not find it important to deworm dogs as a control measure for hydatidosis, though more than half of the participants agreed that deworming livestock and keeping them clean is a true reflection of someone’s social status. There was poor attitude on the disease by participants, which may be contributed to by low literacy level as more than half of them had primary education and below. These barriers related to knowledge and information could hamper the effectiveness of interventions in prevention and control of CH [18]”.

Approximately half of respondents dewormed their livestock; however, more than a quarter of them did not have a regular deworming interval for their livestock. The findings on the number of dog keepers who dewormed their dogs is consistent with a study done in Uganda [16], but was in contrast to studies done in Ethiopia where dog keepers were 71% of the study participants and none of them dewormed their dogs [19] and in Pakistan where dog keepers were 64% and 68% of them dewormed their dogs [7]. The dog owners did not know the risk contained in improper disposal or inappropriate handling of dog faeces in terms of transmission of CH. This makes controlling the disease difficult being that the dog faeces, with infective larvae of the parasite, contaminates the environment hence exposing the livestock and human population to risk of infestation. Control of cystic hydatidosis is less effective without the support of dog-owners, and this support can only be obtained if the people have a clear understanding of the life cycle of the parasite and of risk factors for human and livestock infestations [20].

We noted that slaughtering animals at home was a common practice by respondents, but qualified meat inspectors were rarely contacted to inspect such carcasses. Failure to call a qualified meat inspector to inspect meat slaughtered at home leads not only to improper disposal of infected organs and carcasses, but also risks transmission of other zoonotic diseases to humans. This observation was also made in a study conducted among pastoral communities in Greater Kapoeta of South Sudan [21]. Infested organs and carcasses of cattle slaughtered at home are eaten by the people, fed to dogs, or disposed of in places where dogs can readily access them. Feeding dogs on possibly infested raw meat or organs as done by majority of dog keepers also promote perpetuation of the parasite in dogs and the environment through dog faeces [20]. Our findings revealed that inadequate deworming of dogs and livestock, poor dog faecal disposal and poor disposal of infested organs of animals slaughtered at home are risky practices by Busia communities [19,22].

The prevalence which has been estimated by this study might be lower than expected due to failure to include cattle whose owners could not be traced and therefore not having a chance to establish their infestation status and so the actual risk may be higher than reported. Failure to get positive results on PCR in two occasions might be explained by the fact that using strains to characterize *Echinococcus* is essential to establish a preventive and control strategy in every endemic area, but using DNA based methods for strain/genotype characterizations of *E. granulosus* have some difficulties, especially access to an efficient and pure concentration of DNA and proper primers [23].

There is a non-negligible risk of CH in Busia communities which might worsen with time given that the county is connected to areas perceived to be endemic for the disease (West Pokot and Turkana counties) via livestock trade. The local people also have poor KAP on the disease and hence there is need to implement information, education and communication campaigns to improve KAP on CH in the area. Cystic hydatidosis is an important but neglected zoonotic disease which should be put under surveillance by public health authorities in Kenya. The authors recommend commencement of Busia community public health education (PHE) to improve knowledge, attitude and practices on the disease. The community PHE may also improve veterinary public health activities like deworming dogs, disposing of dog faeces, slaughter hygiene, meat inspection and sanitation measures. Future studies should focus on prevalence of CH in humans and dogs in Busia.

## Acknowledgements

We acknowledge the following institutions and programs for their funding and/or collaboration in this study; Ministry of Health (Kenya Field Epidemiology and Laboratory Training Program), Moi University, International Livestock Research Institute, county governments of Busia and Migori.

We acknowledge the following individuals for their participation or contribution in the development and review of the proposal and/or manuscript; Prof. Eric Fevre, Dr. Dalmas Oyugi, Dr. Mark Nanyingi, Dr. Annie Cook, Dr. Austin Bitek, Dr. Allan Ogendo, Dr. Kelvin Momanyi, Mrs. Mary Midida Owade, Mr. Benard Owade and Mr. Gilbert Nyandiga. We thank Dorothy L Southern for her critical review of the manuscript and her scientific writing support. We also acknowledge the immense contribution made by Kenya Field Epidemiology and Laboratory Training Program (K-FELTP) lecturers and Cohort 12 residents throughout the study period.

## Author Contributions

**Conceptualization:** Henry Joash Ogutu, Bernard Bett, Christina Otieno.

**Data curation:** Henry Joash Ogutu, Bernard Bett, Christina Otieno.

**Formal analysis:** Henry Joash Ogutu, Maurice Owiny.

**Investigation:** Henry Joash Ogutu.

**Methodology:** Henry Joash Ogutu, Maurice Owiny, Bernard Bett, Christina Otieno.

**Software:** Henry Joash Ogutu. Supervision: Bernard Bett, Christina Otieno.

**Supervision:** Bernard Bett, Christina Otieno, Maurice Owiny

**Writing – original draft:** Henry Joash Ogutu, Bernard Bett, Christina Otieno.

**Writing – review & editing:** Henry Joash Ogutu, Maurice Owiny, Bernard Bett, Christina Otieno.

## Conflict of interest

The authors declare that they do not have any competing interest.

## Disclaimer

The findings and conclusions in this manuscript are those of the authors and do not necessarily represent the official position of the Kenyan Ministry of Health or Moi University.

